# AntiCapt: Fine-Tuned Nucleotide Language Models for Predicting and Designing Anticancer Aptamers

**DOI:** 10.64898/2026.09.28.754914

**Authors:** Nisha Bajiya, Naman Kumar Mehta, Gajendra P. S. Raghava

## Abstract

Over the past decade, aptamers have emerged as promising therapeutics, with cancer therapeutics as a major research area. Existing computational methods, however, either predict aptamer-target pairs or design aptamers against a specific target. In this study, we present AntiCapt, a computational method for identifying single-stranded DNA (ssDNA) anticancer aptamers (ACAs) using sequence descriptors and fine-tuned nucleotide language models. The main dataset comprises 1,021 experimentally validated ACAs and an equal number of non-aptamer sequences. Comparative analysis revealed distinct patterns in the nucleotide, dinucleotide, and trinucleotide compositions associated with ACAs. We developed machine learning (ML) models using composition, autocorrelation, binary profiles and structural features. Among these, the best-performing composition-based model achieved an AUC of 0.93 on an independent dataset, while chemical descriptor-and structural feature-based models achieved a maximum AUC of 0.89. ML models using pretrained and fine-tuned NLM embeddings were also developed, with fine-tuned HyenaDNA embeddings achieving the highest performance, with an independent AUC of 0.94. Furthermore, we developed a model for discriminating anticancer aptamers from general aptamers. The best-performing models were integrated into AntiCapt, a web server and a standalone tool for predicting, designing, and genome-scale scanning anticancer aptamers (https://webs.iiitd.edu.in/raghava/anticapt/).

## Introduction

Cancer remains a leading cause of global mortality and continues to pose a major public health challenge despite remarkable advances in diagnosis and treatment.^1,2^ Conventional therapeutic modalities, including surgery, chemotherapy, radiotherapy, immunotherapy, and targeted therapies, have significantly improved overall patient survival.^3,4,5,6^ However, their clinical efficacy is frequently hampered by off-target toxicity, multidrug resistance, tumour heterogeneity, and disease recurrence.^7,8,9,10,11^ Consequently, there is an urgent demand for highly selective molecular recognition agents that can enhance the precision of both cancer diagnosis and therapy.^12,13,14^ Anticancer aptamers (ACAs) have emerged as highly promising therapeutic and diagnostic agents due to their ability to selectively recognize tumour-associated biomarkers and malignant cells.^15,16^ The translational potential of the aptamer drug class is evidenced by two regulatory approvals to date: pegaptanib (Macugen), an anti-VEGF RNA aptamer approved by the US FDA in 2004, and avacincaptad pegol (Izervay), a complement C5-targeting RNA aptamer approved in 2023, both indicated for age-related macular degeneration.^17^

Most notably, the nucleolin-targeting DNA aptamer AS1411 was the first to enter clinical trials in oncology. Furthermore, aptamers targeting key cancer biomarkers, such as epithelial cell adhesion molecule, prostate-specific membrane antigen, and epidermal growth factor receptor, have shown promising preclinical and clinical outcomes.^18,19,20,21,22^ Beyond therapeutic applications, ACAs are widely investigated for targeted drug delivery, cancer diagnostics, biosensors, and molecular imaging.^23,24^ Their structural and chemical adaptability facilitates conjugation with chemotherapeutics, nanoparticles, siRNA, and CRISPR platforms, highlighting their promise for targeted precision medicine.^25,26^

Currently, the identification of functional aptamers depends heavily on the SELEX process, a procedure that is inherently labor-intensive, costly, and time-consuming, frequently requiring multiple iterative selection rounds and empirical validation.^27^ To overcome these limitations, in silico techniques are essential for streamlining aptamer discovery through the rapid computational screening of large sequence spaces.^28,29,30^ Several computational approaches using sequence-derived descriptors and machine learning algorithms have recently been proposed to facilitate aptamer discovery.^31,32^ However, current research focuses almost exclusively on predicting aptamer-target interactions and binding affinities rather than functional disease classes. For instance, AptaNet models aptamer-protein binding via binary classification using k-mer counts and target pseudo-amino acid composition.^33^ More recent architectures, such as AptaTrans using pretrained protein/RNA language encoders,^33^ and AptaBERT employing contrastive/masked language modeling,^34^ further refine target interaction and affinity predictions.

Despite recent advances in aptamer bioinformatics, limited efforts have been made to systematically curate or predict functional therapeutic classes, such as antiviral, antibacterial, or anticancer aptamers. To address this resource gap, we recently established AptViralDB, AptBacterialDB, and AptCancerDB, dedicated repositories compiling experimentally validated antiviral, antibacterial, and anticancer aptamers, respectively. However, while computational prediction tools for therapeutic sequences are well established in related fields, such as the prediction of anticancer peptides (ACPs),^35,36,37,38^ no specialized computational framework currently exists for predicting anticancer aptamers. Developing sequence-based predictive models for anticancer aptamers is therefore essential to bridge this computational gap and accelerate the rational design of aptamer-based cancer therapies.

In this study, we report the first systematic effort to predict anticancer aptamers (ACAs) directly from primary single-stranded DNA sequence data. To accomplish this, we constructed a Main dataset by pairing experimentally validated ACAs from AptCancerDB^39^ with length-matched non-aptamer genomic sequences from NCBI GenBank. To investigate the influence of negative data selection under a biologically stringent benchmark, an Alternate dataset was constructed by pairing the same positive dataset with non-cancer SELEX-derived aptamers compiled from public repositories, including the UTexas Aptamer Database,^40^ AptaBase, and AptaDB.^41^ Ultimately, we integrated our predictive models into AntiCapt, a user-friendly platform designed to enable efficient computational screening, lower the experimental burden of SELEX, and accelerate candidate prioritization for targeted cancer therapy.

## Results

### Primary sequence analysis

#### Nucleotide composition analysis

The average mononucleotide composition (A, T, G, and C), together with the derived AT and GC contents, was calculated for ACAs, general aptamers and random oligonucleotides (Figure 1). ACAs were markedly enriched in guanine relative to random oligonucleotides (28.56% vs. 23.69% mean G content, q < 0.001) and depleted in adenine (22.63% vs. 27.09%, q < 0.001), resulting in a significantly higher GC content in the positive dataset (52.19% vs. 46.81%, q < 0.001) (See Supplementary Table S1.1). These findings are consistent with the well-established tendency of SELEX-derived aptamers, including the G-quadruplex-forming ACAs AS1411, to possess guanine-rich sequences.^42^

**Figure 1.**
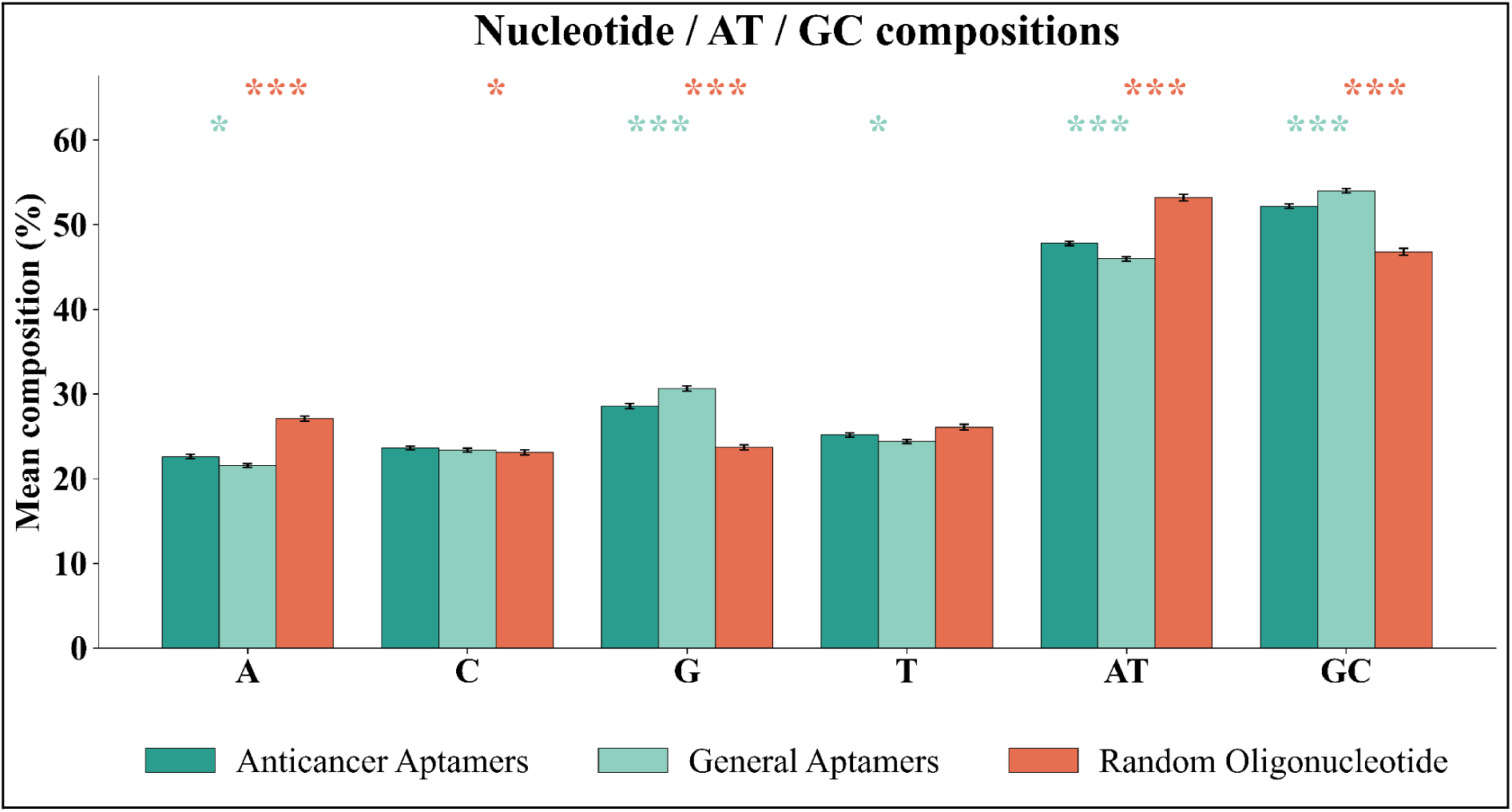
Average nucleotide, AT and GC composition of three nucleotide sets.

In contrast, substantially weaker compositional differences were observed between anticancer and general aptamers. General aptamers exhibited an even higher guanine content than the positive dataset (30.63%, q < 0.001), whereas cytosine composition did not differ significantly (23.63% vs 23.39%, q = 0.57). These observations indicate that generic aptamers already possess many of the nucleotide composition characteristics observed in ACAs.

#### Sequence complexity and homopolymer analysis

Shannon entropy, together with the longest guanine and adenine homopolymer runs, was subsequently compared between datasets (Supplementary Table S1.1). In comparison to random oligonucleotides, ACAs displayed considerable differences in sequence complexity and homopolymer characteristics (q < 0.001). However, these differences largely disappeared when compared with general aptamers, where neither Shannon entropy (q = 0.34) nor the longest adenine homopolymer run (q = 0.96) differed significantly. These results again indicate that sequence complexity and homopolymer organization primarily distinguish ACAs from random oligonucleotides but provide considerably less discrimination from experimentally selected aptamers.^43^

#### Dinucleotide and Trinucleotide Composition

To further determine whether differentiation extended beyond single-nucleotide composition, dinucleotide (16 possible dimers) and trinucleotide (64 possible trimers) frequencies were also compared between datasets. At the dinucleotide level (see Supplementary Table S1.2), 13 of the 16 dinucleotides differed significantly between ACAs and random oligonucleotides, with GG showing the greatest enrichment (8.8% vs. 6.2%) and several adenine/thymine-rich dimers (AA, TA and AT) significantly depleted in the positive dataset, as depicted in Figure 2. In comparison, only nine dinucleotides remained significantly different between anticancer and general aptamers.

**Figure 2.**
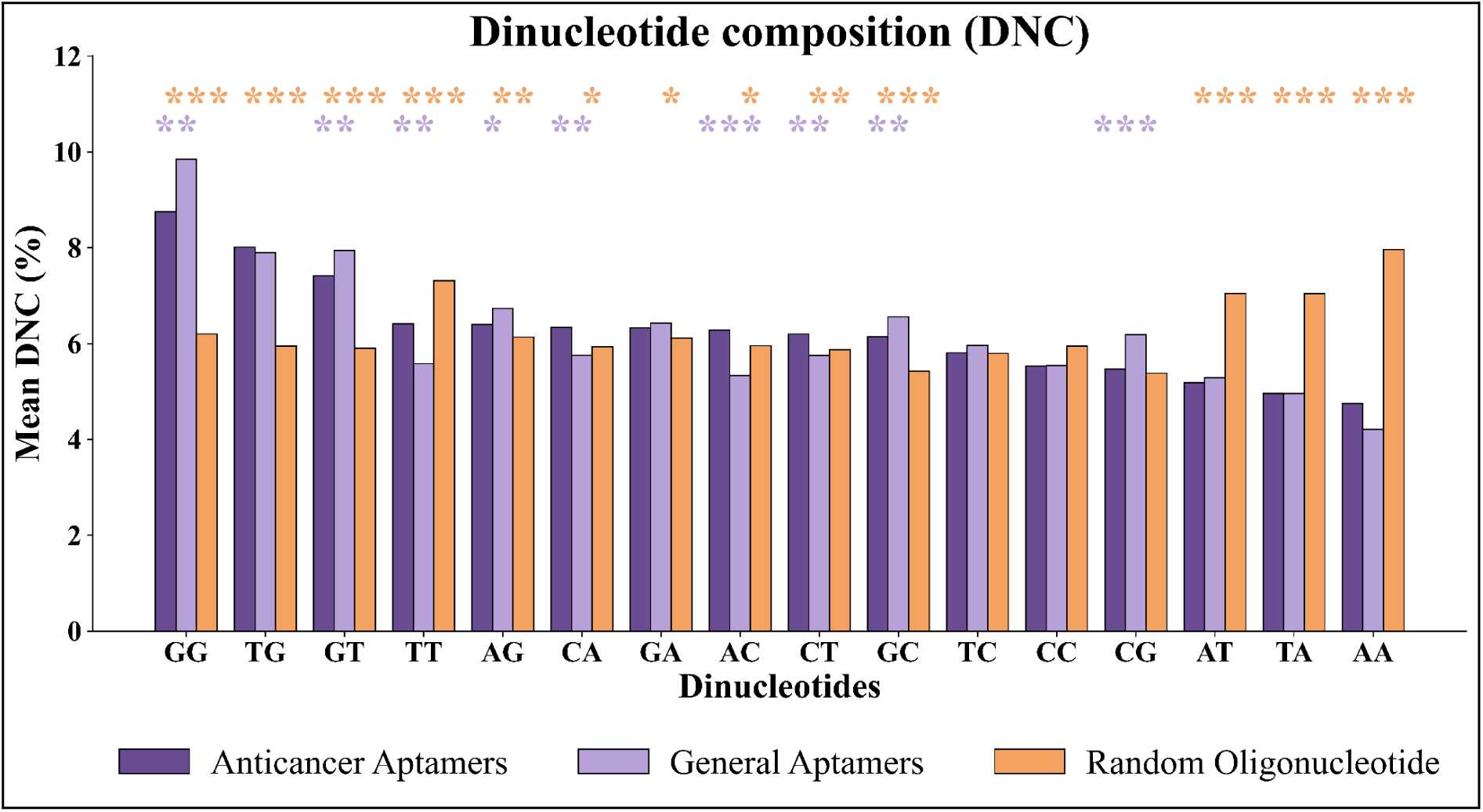
Mean dinucleotide composition (DNC) of Anticancer, General aptamers and Random oligonucleotides (Asterisks denote FDR-corrected significance).

A comparable tendency was also observed for trinucleotide composition, as shown in Figure 3. From the 64 trinucleotides, 41 varied remarkably between ACAs and random oligonucleotides, especially indicating enrichment of G-rich motifs (GGT, GTG, GGG, and TGG) and deficit of A/T-rich motifs (AAA, TAA, ATA and TAT), as summarized in Supplementary Table S1.3. Against general aptamers, only 31 trinucleotides remained significant, and the pattern shifted instinctively. The GGG motif was comparatively lower in ACAs, whereas motifs such as TTT, ACA, and CAA were enhanced. These findings suggest that the distinction between ACAs and general aptamers depends more on specific differences in motif arrangement than on overall guanine enrichment alone.

**Figure 3.**
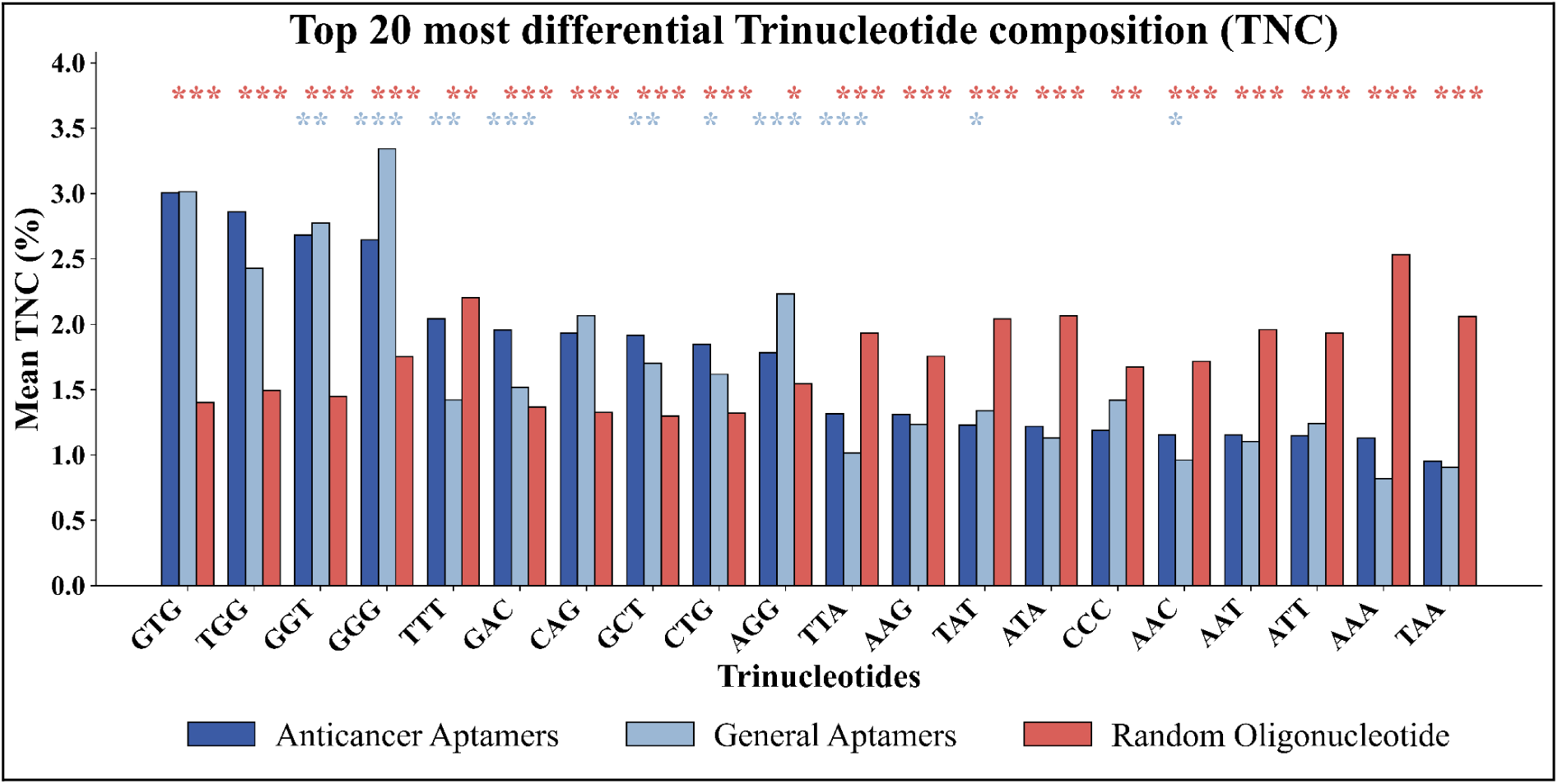
Top 20 FDR-significant trinucleotides between the three sets, ranked by absolute mean composition difference.

#### Secondary structure analysis

The predicted secondary structures of all nucleotide sequences were analyzed using ViennaRNA with DNA-specific nearest-neighbour thermodynamic parameters (Figure 4). Two structural descriptors, including minimum free energy per nucleotide (MFE/nt) and paired fraction, were evaluated. Compared to random oligonucleotides, ACAs were predicted to form particularly more stable and considerably more base-paired secondary structures (with MFE/nt of-0.088 vs.-0.066 kcal/mol/nt at q of 3.5 × 10⁻¹⁵ and paired fraction of 0.43 vs. 0.34 with q of 7.6 × 10⁻²³). These observations are consistent with the SELEX process, which preferentially augments sequences capable of adopting stable folded conformations. The differences were considerably smaller when ACAs were compared with general aptamers. Although both MFE/nt (q = 4.3 × 10⁻³) and paired fraction (q = 2.7 × 10⁻³) remained statistically significant, the mean paired fraction was marginally higher in the general aptamer dataset (0.44 vs 0.43). Together with the primary sequence analyses, these findings indicate that enhanced structural stability is characteristic of functional aptamers generated through SELEX rather than unique to ACAs.

**Figure 4.**
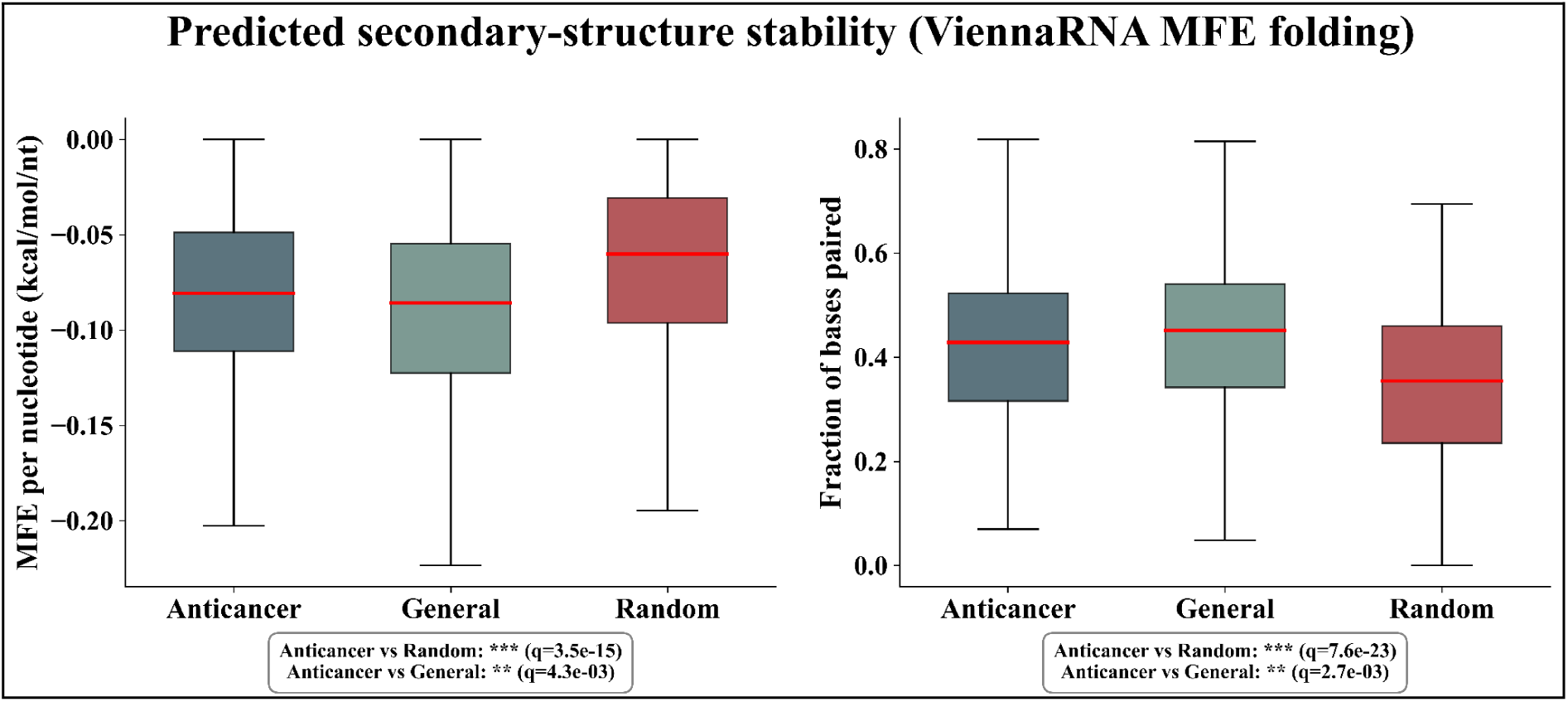
Box plot showing predicted secondary-structure stability (the red line indicates the median).

#### Compositional-based models

The performance of the best ML classifier for each composition-based descriptor family on the main dataset is summarized in Table 1. Among them, the composition family, PKNC, was the strongest with an MCC of 0.756 and AUC of 0.933, outperforming the full concatenated ALLCOMP set (MCC = 0.64 and AUC = 0.919), suggesting that PKNC captures most of the discriminative signal present across the composition descriptor family. The full results are provided in Supplementary Table S2. Also, we have assessed the discriminative power of individual compositional feature columns with the Extra Tree (ET) model (since it’s performing best on the compositional features). In the main dataset, the strongest single composition-derived feature (RDK_AT) from the RDK feature family achieved a test MCC of 0.305 and an AUC of 0.699 (See Supplementary Table S4). These results confirm that no individual descriptor is sufficient for reliable classification and that combining features within and across descriptor families is required. To reduce dimensionality and improve performance, we have applied mRMR feature selection to all compositional features (ALLCOMP). Supplementary Table S6 shows the performance of the ET model on the reduced feature subsets. The Top-100 achieved a validation AUC of 0.916, while the full 123 set achieved an AUC of 0.919, indicating little to no performance improvement.

**Table 1:** Performance of best classifiers on compositional features on validation data of the main dataset.

| Descriptor | Best model | Threshold | Sensitivity | Specificity | MCC | AUC |
| --- | --- | --- | --- | --- | --- | --- |
| **CDK** | ET | 0.49 | 0.828 | 0.771 | 0.6 | 0.886 |
| **DDN** | ET | 0.487 | 0.681 | 0.654 | 0.335 | 0.738 |
| **EN** | RF | 0.497 | 0.745 | 0.624 | 0.372 | 0.765 |
| **ES** | SVM_RBF | 0.51 | 0.691 | 0.551 | 0.245 | 0.679 |
| **NRI** | ET | 0.496 | 0.701 | 0.683 | 0.384 | 0.748 |
| **PDNC** | ET | 0.501 | 0.809 | 0.766 | 0.575 | 0.878 |
| **PKNC** | **ET** | **0.508** | **0.868** | **0.888** | **0.756** | **0.933** |
| **RDK** | LGBM | 0.695 | 0.76 | 0.785 | 0.545 | 0.856 |
| **ALLCOMP** | ET | 0.534 | 0.7794 | 0.8585 | 0.64 | 0.919 |

#### Correlation-based models

Fifteen different types of correlation-based features were evaluated to estimate the performance of ML models in distinguishing ACAs. Among correlation-based descriptors, as shown in Table 2, SC_PTNC achieved the best results on the main datasets, with an MCC of 0.726 and an AUC of 0.941, outperforming the much higher-dimensional ALLCORR concatenation (6,792 features). The complete results are provided in Supplementary Table S3. We have also evaluated the performance of single correlation-derived features on the best-performing ET model (see Supplementary Table S5). The performance on individual features shows that NMBAC_p22_lag1 from the NMBAC family achieved an MCC of 0.307 and an AUC of 0.703, suggesting that individual features are insufficient to differentiate the ACAs and that additional descriptors are required. A similar feature selection approach was also applied to correlation-based feature sets, and the performance of the ET model on the reduced subsets is shown in Supplementary Table S6. The Top-500 correlation features achieved a validation MCC of 0.581 with an AUC of 0.885, whereas 0.905 AUC on the full 6792 set. These observations show that mRMR is effective at reducing dimensionality for the very high-dimensional correlation set and retains additional useful signals beyond the top-ranked features.

**Table 2:** Performance of best classifiers on correlational features on validation data of the main dataset.

| **Descriptor** | **Best model** | **Threshold** | **Sensitivity** | **Specificity** | **MCC** | **AUC** |
| --- | --- | --- | --- | --- | --- | --- |
| **DAC** | XGB | 0.413 | 0.7941 | 0.7756 | 0.5698 | 0.86 |
| **DACC** | SVM_RBF | 0.512 | 0.7108 | 0.9268 | 0.6533 | 0.9168 |
| **DCC** | SVM_RBF | 0.508 | 0.7206 | 0.922 | 0.6562 | 0.9174 |
| **DGAC** | MLP | 0.524 | 0.7059 | 0.7951 | 0.5031 | 0.8474 |
| **DMAC** | MLP | 0.549 | 0.8088 | 0.7561 | 0.5657 | 0.847 |
| **NMBAC** | CBoost | 0.398 | 0.7892 | 0.7024 | 0.4935 | 0.8575 |
| **PC_PDNC** | ET | 0.51 | 0.799 | 0.8098 | 0.6088 | 0.8823 |
| **PC_PTNC** | ET | 0.492 | 0.8676 | 0.8634 | 0.7311 | 0.9363 |
| **SC_PDNC** | ET | 0.47 | 0.8333 | 0.7659 | 0.6005 | 0.8827 |
| **SC_PTNC** | **ET** | **0.499** | **0.8529** | **0.8732** | **0.7263** | **0.9408** |
| **TAC** | ET | 0.547 | 0.6225 | 0.8293 | 0.4619 | 0.7926 |
| **TACC** | SVM_RBF | 0.562 | 0.7304 | 0.7902 | 0.5216 | 0.8372 |
| **TCC** | SVM_RBF | 0.539 | 0.75 | 0.7561 | 0.5061 | 0.8359 |
| **TGAC** | SVM_RBF | 0.477 | 0.7941 | 0.6878 | 0.4846 | 0.8244 |
| **TMAC** | XGB | 0.546 | 0.7059 | 0.7805 | 0.4878 | 0.8161 |
| **ALLCORR** | LGBM | 0.462 | 0.848 | 0.8049 | 0.6535 | 0.9074 |

#### Binary profile-based models

We have also evaluated binary positional encodings to distinguish the ACAs from the random oligonucleotides. As shown in Table 3, binary encodings performed competitively despite their very high dimensionality. On the main Dataset, the combined mono/di-position encoding (BPM) with ET yielded the best result (MCC of 0.707 and AUC of 0.929), closely followed by ALLBINARY with GPU-accelerated CBoost (AUC of 0.926 and MCC of 0.687). The completed binary results for the main dataset are provided in Supplementary Table S7.

**Table 3:** Performance of best classifiers on binary features on the validation data of the main dataset.

| Descriptor | Best model | Accuracy | Sensitivity | Specificity | MCC | AUC |
| --- | --- | --- | --- | --- | --- | --- |
| BP_DP | CBoost_GPU | 0.841 | 0.819 | 0.863 | 0.683 | 0.920 |
| BP_TP | LR | 0.829 | 0.828 | 0.829 | 0.658 | 0.912 |
| BPM | ET | **0.848** | **0.765** | **0.932** | **0.707** | **0.929** |
| BPD | ET | 0.834 | 0.750 | 0.917 | 0.677 | 0.918 |
| BPT | MLP | 0.831 | 0.794 | 0.868 | 0.664 | 0.917 |
| ALLBINARY | CBoost_GPU | 0.844 | 0.828 | 0.859 | 0.687 | 0.926 |

#### Structural (secondary-structure and shape) features-based models

The combined secondary-structure and DNA-shape feature vector was evaluated across 18 different types of classifiers, and their performance is shown in Table 4, with complete results provided in Supplementary Table S8. MLP gave the strongest result with a validation accuracy of 0.809, an MCC of 0.619, and an AUC of 0.891, with HistGradientBoosting a close second (MCC = 0.614, AUC = 0.883). Structural features, therefore, carry a real, moderate discriminative signal, consistent with the secondary-structure differences as described in primary analysis. However, their performance is no better than that of the compositional descriptors or the fine-tuned HyenaDNA.

**Table 4:** Performance of ML models on predicted structural features on the validation data of the main dataset.

| Model | Accuracy | Sensitivity | Specificity | MCC | AUC |
| --- | --- | --- | --- | --- | --- |
| MLP | **0.809** | **0.784** | **0.834** | **0.619** | **0.891** |
| HistGB | 0.807 | 0.828 | 0.785 | 0.614 | 0.883 |
| LGBM | 0.819 | 0.853 | 0.785 | 0.640 | 0.883 |
| ET | 0.790 | 0.809 | 0.771 | 0.580 | 0.875 |
| XGB | 0.782 | 0.779 | 0.785 | 0.565 | 0.870 |
| CBoost | 0.782 | 0.789 | 0.776 | 0.565 | 0.862 |
| SVM_RBF | 0.790 | 0.799 | 0.781 | 0.580 | 0.859 |
| RF | 0.787 | 0.779 | 0.795 | 0.575 | 0.855 |
| Bagging | 0.775 | 0.784 | 0.766 | 0.550 | 0.853 |
| GBoost | 0.753 | 0.765 | 0.742 | 0.506 | 0.837 |
| KNN | 0.765 | 0.828 | 0.702 | 0.535 | 0.836 |
| LR | 0.712 | 0.716 | 0.707 | 0.423 | 0.795 |
| SVM_Linear | 0.719 | 0.716 | 0.722 | 0.438 | 0.794 |
| **LDA** | 0.716 | 0.721 | 0.712 | 0.433 | 0.792 |
| **AdaB** | 0.704 | 0.760 | 0.649 | 0.411 | 0.783 |
| **NB** | 0.665 | 0.593 | 0.737 | 0.333 | 0.759 |
| **QDA** | 0.516 | 0.034 | 0.995 | 0.106 | 0.758 |
| **DT** | 0.709 | 0.696 | 0.722 | 0.418 | 0.709 |

#### Chemical descriptors-based models

Table 5 shows the performance of the best models on the SMILE-based features on the main dataset. Among the RDKit scalar descriptor sets, the full 0D+1D+2D combination (201 features) and the 2D subset alone (90 features) performed almost identically and yielded the strongest results. CatBoost on the 2D subset reached validation MCC of 0.614 with an AUC of 0.891, and Gradient Boosting on the full combined set reached validation MCC of 0.653 and an AUC of 0.892. While the structural fingerprints (Morgan, MACCS, RDKitFP, and Layered) showed extremely poor performance, as summarized in Supplementary Table S9. Every classifier evaluated on each of these fingerprints returned a validation MCC indistinguishable from zero, indicating essentially no discriminative signal. While the AtomComposition, where XGBoost reached a validation MCC of 0.472 and an AUC of 0.826, is comparable to RDKit’s 1D descriptor subset. The lower performance of fingerprints such as Morgan and MACCS is because they are designed to detect the aromatic rings, functional groups, and heterocyclic scaffolds that distinguish diverse drug-like small molecules from one another, whereas aptamers are built from the same sugar-phosphate DNA backbone and only four base types, making them hard to distinguish from the random oligonucleotides.

**Table 5:**
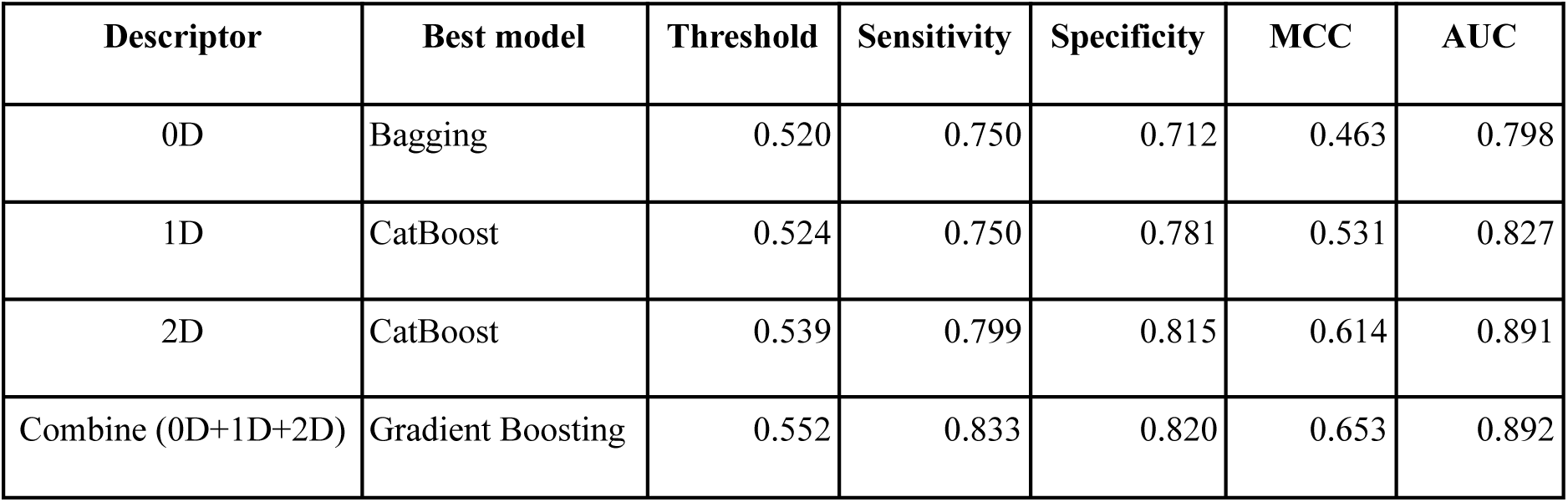
Performance of best classifier on chemical descriptors on the validation data of the main dataset.

| Descriptor | Best model | Threshold | Sensitivity | Specificity | MCC | AUC |
| --- | --- | --- | --- | --- | --- | --- |
| 0D | Bagging | 0.520 | 0.750 | 0.712 | 0.463 | 0.798 |
| 1D | CatBoost | 0.524 | 0.750 | 0.781 | 0.531 | 0.827 |
| 2D | CatBoost | 0.539 | 0.799 | 0.815 | 0.614 | 0.891 |
| Combine (0D+1D+2D) | Gradient Boosting | 0.552 | 0.833 | 0.820 | 0.653 | 0.892 |

#### Nucleotide language embeddings-based models

Nucleotide-based large language models and pretrained embeddings have been widely used for various tasks for genomic sequences.^44,45^ Table 6 compares pretrained and fine-tuned embeddings from the three NLMs. Fine-tuning substantially and consistently improved performance across all three models. The fine-tuned HyenaDNA embeddings with an LGBM classifier achieved the best language-model result, achieving an MCC of 0.726 and an AUC of 0.944. Pretrained embeddings without fine-tuning were markedly weaker than their fine-tuned counterparts, indicating that generic NLM representations require task-specific adaptation to capture anticancer-aptamer-relevant signals. The complete performance is summarized in Supplementary Table S10.

**Table 6:** Performance of best classifier on pretrained vs. fine-tuned NLM embeddings on the validation data of the main dataset.

| Embedding model | Best model | Accuracy | Sensitivity | Specificity | MCC | AUC |
| --- | --- | --- | --- | --- | --- | --- |
| **Pretrained Embeddings** |  |  |  |  |  |  |
| **HyenaDNA** | MLP | 0.770 | 0.760 | 0.780 | 0.540 | 0.869 |
| **DNABERT (6-mer)** | SVM_RBF | 0.814 | 0.799 | 0.829 | 0.629 | 0.893 |
| **Nucleotide Transformer (500M)** | MLP | 0.719 | 0.691 | 0.746 | 0.438 | 0.817 |
| **Finetuned Embeddings** |  |  |  |  |  |  |
| **HyenaDNA** | **LGBM** | **0.863** | **0.858** | **0.868** | **0.726** | **0.944** |
| **DNABERT (6-mer)** | CBoost | 0.858 | 0.828 | 0.888 | 0.718 | 0.933 |
| **Nucleotide Transformer (500M)** | LR | 0.851 | 0.824 | 0.878 | 0.703 | 0.924 |

#### Discrimination of Anticancer aptamers from general aptamers

Another dataset was constructed in this study, called the alternate dataset, which consists of 1,021 ACAs and an equal number of experimentally validated non-cancer general aptamers, rather than a random genomic background. Based on the above results, the LGBM with fine-tuned HyenaDNA embeddings achieved the best performance and was selected as the final model. We have utilized this framework to evaluate the performance on the validation data of the alternate dataset. Here, the HyenaDNA was fine-tuned with the training set of the alternate dataset, and the performance of ML models is summarized in Table 7 and the complete results in Supplementary Table S11. Across all ML models, the ET model showed the highest performance, with an MCC of 0.594 and an AUC of 0.891. However, performance on the alternate dataset is lower than on the main dataset, reflecting a genuine, intrinsic difficulty in distinguishing cancer-targeting aptamers from other functional aptamers, as general aptamers already share many compositional and structural signatures with ACAs.

**Table 7:** Performance of the ML models on fine-tuned HyenaDNA embeddings on validation data on the alternate dataset.

| **Models** | **Accuracy** | **Sensitivity** | **Specificity** | **Precision** | **F1** | **MCC** | **AUC** |
| --- | --- | --- | --- | --- | --- | --- | --- |
| **ET** | 0.797 | 0.805 | 0.789 | 0.793 | 0.799 | 0.594 | 0.891 |
| **RF** | 0.790 | 0.795 | 0.784 | 0.787 | 0.791 | 0.579 | 0.882 |
| **XGB** | 0.807 | 0.820 | 0.794 | 0.800 | 0.810 | 0.614 | 0.881 |
| **MLP** | 0.804 | 0.810 | 0.799 | 0.802 | 0.806 | 0.609 | 0.880 |
| **AdaB** | 0.792 | 0.771 | 0.814 | 0.806 | 0.788 | 0.585 | 0.878 |
| **LGBM** | 0.807 | 0.824 | 0.789 | 0.797 | 0.811 | 0.614 | 0.878 |
| **LR** | 0.804 | 0.820 | 0.789 | 0.796 | 0.808 | 0.609 | 0.876 |
| **CBoost** | 0.795 | 0.790 | 0.799 | 0.798 | 0.794 | 0.589 | 0.874 |
| **GBoost** | 0.807 | 0.820 | 0.794 | 0.800 | 0.810 | 0.614 | 0.871 |
| **SVM_Linear** | 0.782 | 0.776 | 0.789 | 0.787 | 0.781 | 0.565 | 0.866 |
| **SVM_RBF** | 0.792 | 0.785 | 0.799 | 0.797 | 0.791 | 0.584 | 0.864 |
| **NB** | 0.795 | 0.805 | 0.784 | 0.789 | 0.797 | 0.589 | 0.845 |
| **KNN** | 0.802 | 0.795 | 0.809 | 0.807 | 0.801 | 0.604 | 0.836 |
| **DT** | 0.760 | 0.761 | 0.760 | 0.761 | 0.761 | 0.521 | 0.760 |

#### Architecture of the AntiCapt web server

To predict ACAs, a web-based service called “AntiCapt” was implemented, incorporating the best-performing fine-tuned HyenaDNA models as selected for the main dataset. The web server consists of three major modules: Predict, Design and Scan. The *Predict* module allows users to provide an input query nucleotide sequence in a single or multiple FASTA files, compute its fine-tuned HyenaDNA embedding, and return a prediction score, GC content, and an anticancer/non-anticancer classification using the main dataset’s final model. If users are aware that their input sequences are aptamers, then they can opt for an alternate dataset that allows classification of anticancer aptamers from general non-cancer aptamers. The *Design* module generates an independent point mutant from the input nucleotide sequence and predicts the anticancer potential of the mutant variants using the main dataset embedding model, enabling a user to identify which substitutions are predicted to increase or decrease anticancer potential without regenerating a hand-crafted descriptor vector for each variant. The *Scan* module utilizes a sliding window approach across a longer candidate nucleotide sequence or aptamer library, flagging the regions or sub-regions with high predicted anticancer potential using the default main dataset model. In addition, the standalone package, the GitHub repository (https://github.com/raghavagps/AntiCapt), and the PyPI package (“pip install anticapt”) are also available in the *Download* module, along with the training and validation sets.

## Discussion and Conclusion

The present study develops an LLM-based framework to identify, design, and scan anticancer aptamers directly from nucleotide sequences. We demonstrate that ACAs exhibit distinct sequence patterns that enable reliable computational classification by rigorously analysing several sequence descriptors, feature selection techniques, traditional ML algorithms, and pretrained or fine-tuned NLMs. Aside from developing a predictive model, this study provides biological and methodological insights into the sequence properties of ACAs, emphasising the need for physiologically meaningful benchmark datasets to assess computational prediction methods.

The primary sequence investigated in the study indicates that ACAs exhibit unique compositional and structural properties compared with random oligonucleotide sequences. ACAs were higher in guanine and GC content, had distinctive di-and trinucleotide patterns, and were predicted to have more stable secondary structures than random oligonucleotides. These findings are consistent with the known structural requirements of functional aptamers, which indicate that high GC content promotes the formation of stable secondary structures that enable high-affinity target recognition, such as stem-loops and G-rich conformations. Guanine-rich sites seen in several experimentally confirmed ACAs, such as the well-studied AS1411 aptamer, can create G-quadruplex structures that support molecular recognition, stability, and biological activity.^46^ All of these results point to sequence composition and structural stability as key factors that determine aptamer functionality.

However, a comparison to experimentally verified non-cancer aptamers revealed a more subtle biological picture. When ACAs were compared to generic aptamers, many of the sequence properties that separated them from random oligonucleotides were far less evident. Differences in GC content, sequence complexity, homopolymer organization, and projected folding stability were significantly decreased. This suggests that many of these traits are generic aspects obtained throughout the SELEX process rather than anticancer-specific characteristics. Instead, differentiation between anticancer and non-cancer aptamers appeared to be predominantly based on minor changes in local sequence organization, motif placement, and higher-order sequence context. As a result, structural characteristics such as G-richness and increased foldability should be seen as general hallmarks of functional aptamers rather than unique indicators of anticancer activity.

The performance of the various feature representations adds to this biological understanding. Correlation-based sequence representations, namely PseDNC and PseTNC, consistently outperformed basic nucleotide composition descriptors on the main dataset. Compared with composition-based features, these descriptors preserve short-range sequence order while retaining physical interactions between nearby nucleotides. Their superior performance suggests that local sequence structure has a stronger impact on anticancer aptamer identification than nucleotide frequencies alone. Notably, fusing many features into large concatenated feature sets did not always improve prediction accuracy, suggesting that indiscriminate feature expansion introduces redundancy and noise that can mask valuable sequence patterns. This conclusion is further supported by the mRMR feature-selection analysis, in which relatively small subsets of informative features achieved performance comparable to or exceeding that of substantially larger feature sets.

Despite the lack of precise physicochemical information and the creation of exceptionally high-dimensional feature spaces, binary positional encodings demonstrated remarkable predictive ability. Another finding from this study concerns the performance of NLMs embeddings. Although genomic language models like HyenaDNA, DNABERT, and the Nucleotide Transformer have performed admirably across a variety of genomic prediction tasks, their pretrained embeddings have underperformed compared with engineered sequence descriptors for ACA prediction. Only after task-specific fine-tuning did these models become competitive with conventional machine learning approaches. This finding is likely due to the significant variations between the genomic sequences utilized in large-scale pretraining and the comparatively limited, structurally constrained SELEX-derived sequence space occupied by aptamers. As a result, broad genomic representations do not simply translate to aptamer prediction without additional domain adaptation. Also, in the alternate dataset, performance with experimentally validated non-cancer aptamers provides a considerably more challenging and biologically realistic benchmark because they already possess many of the structural and compositional characteristics shared by functional aptamers.

From a practical perspective, AntiCapt provides an efficient computational framework for categorizing ACA candidates before experimental validation. Since the model requires only the primary nucleotide sequence as input, it can rapidly screen large candidate libraries generated during SELEX and identify promising sequences for downstream experimental characterization. Such computational pre-screening has the potential to substantially reduce the experimental burden, cost, and time associated with iterative aptamer selection while increasing the efficiency of therapeutic aptamer discovery. AntiCapt is expected to facilitate the rapid discovery and development of aptamer-based cancer therapeutics.

## Materials and Methods

The overall AntiCapt workflow, from data curation through feature generation, feature selection, model benchmarking and final tool deployment, is summarized in Figure 5.

**Figure 5.**
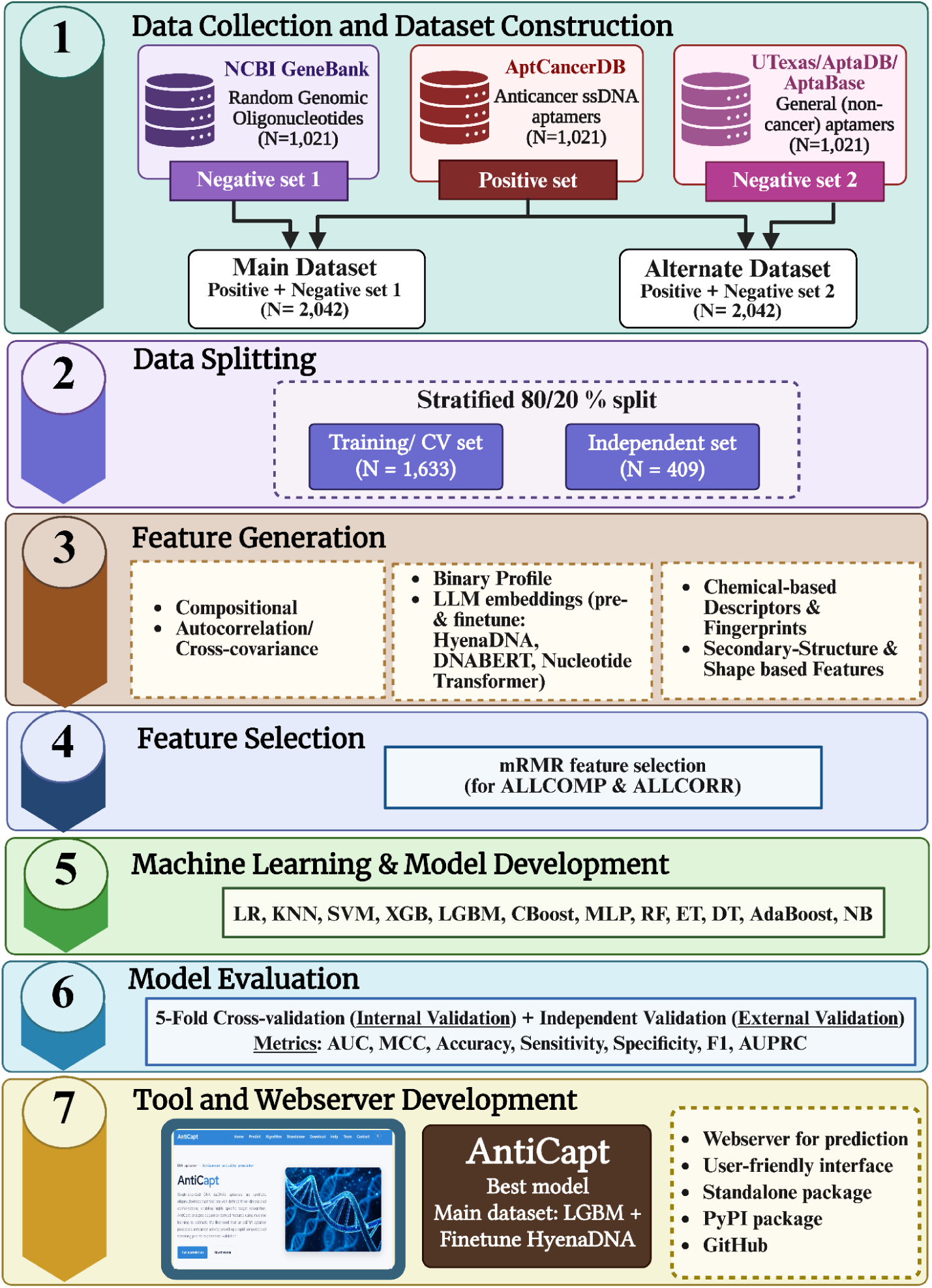
Overall AntiCapt workflow.

### Datasets construction

In this study, we have created two datasets: the main and the alternate. Anticancer aptamers (ACAs) were compiled from AptCancerDB, a database of DNA and RNA aptamers reported against cancer-associated protein, cell and glycolipid targets.^39^ Since the majority of data entries are ssDNA, we have selected only ssDNA aptamers. After removing modified, duplicate, and non-natural nucleotides and ambiguous entries, 1,021 unique, non-redundant anticancer DNA aptamers were obtained, ranging from 13 to 118 nucleotides in length (mean 65.6 nt). These 1021 ACAs were used for creating the following two datasets;

### For Anticancer aptamers versus Random oligonucleotides (Main dataset)

Our main dataset contains ACAs as positive sequences and random oligonucleotides as negative sequences. We obtained an equal number of length-matched sequences from the NCBI GenBank nucleotide repository (13-120 nt, mean 66.4 nt) that are not known aptamers and were used to represent a generic, non-selected nucleic acid background, thereby providing a compositionally distinct negative class. In summary, our main dataset contains 1,021 experimentally validated ACAs and 1,021 non-aptameric, oligonucleotide sequences.

### For Anticancer aptamers versus General aptamers (Alternate dataset)

Our alternate dataset includes anticancer and non-cancer aptamers. To obtain non-cancer aptamers, we compiled sequences from general-purpose, non-cancer-specific aptamer repositories, namely the UTexas Aptamer Database,^40^ AptaDB,^41^ and AptaBase (https://www.iitg.ac.in/proj/aptabase/about.html). These are known aptamers selected by SELEX against non-cancer targets (e.g., small molecules, inhibitors, viral proteins, non-oncology human proteins) and constitute a biologically realistic negative class that shares the general sequence and structural characteristics of aptamers with the positive set. To make a balanced dataset, we randomly selected 1,021 length-matched aptamer sequences (13-117 nt, mean 66.4 nt) from this pool that had no reported association with cancer. In simple terms, our alternative dataset contains 1,021 ACAs and 1,021 non-cancer aptamer sequences.

### Primary sequence-level analysis

To explore the inherent capability of the ACA sequence before ML model development, a comprehensive primary sequence-level analysis was conducted on the positive dataset (ACAs) and the two independently constructed negative datasets (Random Oligonucleotides and General Aptamers). The analyses were designed to identify compositional and structural differences between ACAs and their respective negative backgrounds.

### Nucleotide composition (NC) analysis

Mononucleotide composition (MNC), dinucleotide composition (DNC), and trinucleotide composition (TNC) were calculated for every sequence. MNC was computed as the percentage occurrence of each nucleotide (A, C, G, and T). At the same time, DNC and TNC were calculated as the percentage frequency of overlapping di-and trinucleotide k-mers, respectively. In addition, the overall adenine-thymine (AT) content and guanine-cytosine (GC) content were determined for each sequence. The equations 1 to 4 are as follows;

#### Mono-nucleotide composition (%)

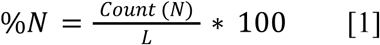

#### AT / GC content

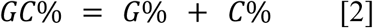

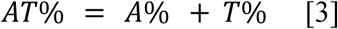

#### k-mer (di-/tri-nucleotide) composition (%)

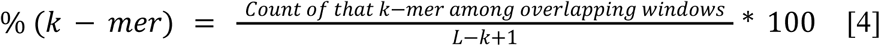

Where, N = A, T, G, C and L = sequence length

#### Sequence complexity and homopolymer analysis

The Shannon entropy of the mononucleotide distribution was utilized to quantify the sequence complexity.^47^ It is expressed in bits, where a value of 2.0 represents a perfectly uniform nucleotide composition, as shown in equation 5. Also, to evaluate sequence patterns associated with structural motifs, the longest continuous guanine (G) and adenine (A) homopolymer runs were identified for each sequence as proxies for G-and A-rich sequence tracts.

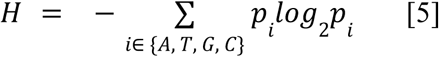

Where, *p_i_* = frequency of base i

#### Secondary structure prediction

To complement the primary sequence analyses, the secondary structure of every DNA aptamer was predicted using the ViennaRNA package 2.0 with the DNA-specific nearest-neighbour thermodynamic parameter set described by Mathews et al. rather than the default RNA thermodynamic parameters.^48^ Since the thermodynamic properties governing DNA duplex formation differ substantially from those governing RNA duplex formation, DNA-specific parameters were selected.^49^ For each aptamer sequence, two structural descriptors were calculated, (i) Minimum free energy per nucleotide (MFE/nt) (in kcal/mol), which represents the thermodynamic stability of the predicted secondary structure (where lower values indicate greater structural stability), and (ii) Paired fraction, which is defined as the proportion of nucleotides predicted to participate in canonical Watson-Crick base pairing.

#### Statistical analysis

For all continuous sequence-derived features, including nucleotide compositions, AT/GC content, Shannon entropy, and homopolymer lengths and secondary structure, ACAs were compared independently with each negative dataset using the two-sided Mann-Whitney U test.^50^ This non-parametric test was selected because these variables are not assured to follow a normal distribution and because it compares the relative distributions of two independent groups without assuming a specific underlying distribution. Since multiple sequence features were evaluated simultaneously, the resulting p-values were adjusted using the Benjamini-Hochberg false discovery rate (FDR) procedure,^51^ and statistical significance was assessed at an FDR-adjusted *q*-value < 0.05. The magnitude of the observed differences was quantified using the rank-biserial correlation, which was calculated for each feature using the Mann-Whitney U statistic.^52^ For the binary G-quadruplex motif feature, differences in motif occurrence between groups were evaluated using Fisher’s exact test, and the corresponding odds ratio was reported.^53^

#### Sequence-based Feature Generation

To develop ML models that can classify ACAs from the non-ACAs, various sequence-based features were computed using Nfeature, an in-house platform for generating composition-, distance-distribution-, correlation-and binary-profile-based features of nucleotide sequences.^54^

#### Composition-based descriptors

Using the Composition module of Nfeature, eight composition-based descriptors were computed as shown in Supplementary Table S12.1. These are nucleotide/di-nucleotide k-mer composition (CDK), distance distribution of nucleotides (DDN), sequence-level entropy (EN), nucleotide-level entropy (ES), nucleotide repeat index (NRI), pseudo-dinucleotide composition (PDNC), pseudo k-tuple nucleotide composition (PKNC) and reverse-complement dinucleotide k-mer composition (RDK). These eight sub-families were concatenated into a combined 123-dimensional composition feature set (ALLCOMP).

#### Correlation-based descriptors

Similar to compositional features, Nfeature was also employed to compute various auto-and cross-covariance descriptors using the Correlation module to develop ML models. Fifteen descriptor sub-families that capture sequence order and physicochemical correlations were generated, as summarized in Supplementary Table S12.2. Dinucleotide-based auto-covariance (DAC), dinucleotide-based auto-cross-covariance (DACC), dinucleotide-based cross-covariance (DCC), dinucleotide-based Geary autocorrelation (DGAC), dinucleotide-based Moran autocorrelation (DMAC), normalized Moreau-Broto autocorrelation (NMBAC), parallel-and series-correlation pseudo-dinucleotide and pseudo-trinucleotide composition (PC-PseDNC/PC_PDNC, SC-PseDNC/SC_PDNC, PC-PseTNC/PC_PTNC, SC-PseTNC/SC_PTNC), and their trinucleotide auto/cross-covariance counterparts (TAC, TACC, TCC, TGAC, TMAC) were among them. All fifteen sub-families were concatenated into a combined 6,792-dimensional correlation feature set named ALLCORR.

#### Binary-based encodings

Nfeature was also used to generate five types of binary/one-hot positional encoding schemes via the Binary Profile module. As shown in Supplementary Table S12.3, it includes mono-and di-nucleotide binary position encoding (BP_DP), tri-nucleotide binary position encoding (BP_TP), and binary encodings of mono-(BPM), di-(BPD) and tri-nucleotide (BPT) content, concatenated into an ALLBINARY vector whose dimensionality scales with the maximum sequence length.

#### Chemical-Based Molecular Descriptors and Fingerprints

In addition to the sequence-derived descriptors and LLM embeddings, each aptamer was also represented as a small-molecule-style feature vector. This was done by first converting aptamer sequences (5′ to 3′ ssDNA) into the corresponding oligonucleotide SMILES strings, explicitly capturing the sugar-phosphate backbone and base composition, using a custom RDKit-based builder.^55^ After computing RDKit molecular descriptors, structural fingerprints, and atom-composition features from these SMILES representations, a total of 10,637 features per molecule were obtained.

RDKit’s descriptor calculator was used to compute 201 physicochemical and topological descriptors (excluding 16 computationally unstable descriptors that require graph eigenvalue decomposition). These were further classified by dimensionality as follows;

#### 0D Descriptors

Generated 26 feature vectors of constitutional/counting descriptors that summarize the molecule as a whole, independent of its connectivity, such as molecular weight and atom, bond, and ring counts.

#### 1D Descriptors (Fingerprints and Structural Keys)

It consists of 85 feature vectors that describe fragment/substructure counts within the molecule, most commonly represented as molecular fingerprints. Seven-bit-or count-based fingerprint and composition representations were computed, as summarized in Supplementary Table S12.4. Six RDKit fingerprint types, including Morgan, MACCS, RDKit topological, Topological Torsion, Pattern, and Layered, were computed (2,048 bits each, except MACCS at 167 bits), totalling 10,407 fingerprint features.

#### 2D Descriptors

It consists of 90 feature vectors that describe the molecular topology based on the graph representation of the molecule, such as the number of rotatable bonds and rings, topological polar surface area, Crippen LogP, Labute approximate surface area, and the fraction of sp3-hybridized atoms. Additionally, 29 atom-composition features were computed directly from the molecular graph, capturing the relative frequency of individual elements (monoatom), bonded element pairs (diatom), and bonded element triples (triatom).

#### Secondary-Structure and Shape-Based Features

To complement the above features, we have also computed a feature set of 29 vector-sized features directly from each aptamer’s predicted secondary structure and sequence-dependent local DNA structure, combining composition-and folding-based descriptors. These includes the GC_content, AT_content, Sequence_length, the sixteen possible dinucleotide compositions (Dinuc_AA through Dinuc_CC), MFE (from the ViennaRNA-predicted secondary structure), Hairpin_stems, Max_stem_length, and Max_loop_length (extracted from the corresponding dot-bracket notation), a G4_score quantifying G-quadruplex-forming propensity, and DNA_bendability, a sequence-dependent flexibility parameter. Four additional local DNA shape features, which are minor groove width (MGW), Roll, propeller twist (ProT), and helix twist (HelT), were also incorporated using the pentamer-based, sliding-window DNAshape model, which predicts these structural parameters directly from nucleotide sequence without requiring a solved structure.^56^ Since several of these features (MGW, Roll, ProT, HelT) are defined at each nucleotide position along the sequence, sequence-averaged summary statistics were used to produce a fixed-length feature vector that is comparable across aptamers of different lengths.

#### Nucleotide Language model (NLM) embeddings

Apart from conventional descriptors, sequence embeddings were also generated using three pretrained nucleotide (DNA) language models, which are DNABERT (6-mer tokenisation, DNA_bert_6), HyenaDNA, and the Nucleotide Transformer, that differ considerably in architecture, tokenization strategy, and the type of sequence dependency, as summarized in Supplementary Table S12.5. DNABERT, based on the BERT bidirectional transformer encoder for genomic sequences, tokenizes DNA into overlapping 6-mers and trains it via masked k-mer prediction. Local sequence motifs, including regulatory elements and promoter-like signals, were captured very well using this design, but, similar to other standard transformer encoders, complexity scales quadratically with sequence length, limiting the practical context.^57^ HyenaDNA employs the Hyena operator, which is an attention-free architecture built from implicit long convolutions and gated operators, tokenizing sequences at the level of single nucleotides. Since its computational cost scales sub-quadratically with sequence length, HyenaDNA can be trained on context windows spanning hundreds of thousands to over a million nucleotides in its larger configurations. Orders of magnitude longer than a standard transformer encoder could practically support, making it particularly well-suited to modelling long-range dependencies relevant to a folded aptamer structure spanning its full sequence length.^58^ The Nucleotide Transformer, developed by InstaDeep, is the genomic counterpart to large protein language models such as ESM2 and ProtT5. It is a transformer encoder trained via masked k-mer prediction on billions of nucleotides drawn from the human reference genome, an additional 3,202 diverse human genomes, and 850 genomes spanning a wide range of species, giving it broad exposure to genomic sequence context and functional regions beyond what a single-genome or single-species training corpus would provide.^59^ Both pretrained embeddings and those fine-tuned on the AntiCapt training set were deployed across all three LLM models to assess whether task-specific transformation improves over generic pretrained representations.

#### Feature selection

To reduce dimensionality and redundancy in the composition (123 features) and correlation (6,792 features) features, Minimum Redundancy Maximum Relevance (mRMR) feature selection was applied.^60^ It contains the relevant feature sets with reduced similarity, and classifiers were retrained on the top-K ranked features for a range of K values (Composition = 10,20,30,50,70,90 & 100; Correlation = 100, 200, 300, 400, 500, 700, 900 & 1000), allowing direct comparison of predictive performance against the full, unreduced feature set.

#### Model development

To develop classification models, various types of ML algorithms have been evaluated in this study. The classifiers span linear, instance-based, kernel, neural and tree-ensemble methods on every generated feature set. These are Logistic Regression (LR), k-Nearest Neighbours (KNN), Gaussian Naive Bayes (NB), Decision Tree (DT), Support Vector Machines with linear and radial basis function kernels (SVM_Linear, SVM_RBF), Multi-Layer Perceptron (MLP), Random Forest (RF), Extra(mely Randomized) Trees (ET), AdaBoost (AdaB), Gradient Boosting (GBoost), XGBoost (XGB), LightGBM (LGBM) and CatBoost (CBoost). In case of high-dimensional binary encodings, GPU-accelerated CatBoost, XGBoost, and LightGBM implementations were utilized to improve computational tractability. All models were run with default hyperparameters as a baseline, and a subset of the strongest feature sets was then optimized via hyperparameter search to assess the marginal benefit of tuning.

#### Cross-validation and performance evaluation

To provide internal and external validation, we divided each of our datasets into training and validation sets in an 80-20% ratio (1,633 and 409 sequences). For internal validation, we employed the standard five-fold CV technique, and model and hyperparameter selection were performed entirely on this internal five-fold CV training set. To validate the performance of our models on unseen data, we evaluated the selected models on the 20% validation set, which was not used at any stage of training or model selection; this is called external validation.^33,61^

We have utilized both threshold-dependent and-independent parameters to evaluate the performance of the developed models, as explained in previous studies.^62^ Sensitivity, specificity, accuracy, and the Matthews correlation coefficient (MCC), as representative of threshold-dependent parameters, were computed. While to assess threshold-independent performance, the area under the curve (AUC) was used for model comparison and ranking, as it remains informative in the presence of class overlap and does not depend on a single classification threshold. MCC was employed as the primary ranking metric because it remains informative under class overlap and is considered more robust than accuracy or F1 for balanced binary classification benchmarking.^63^

### Limitations and future prospects

There are several limitations of this study which should be acknowledged. First, the positive set is limited to AptCancerDB-annotated DNA aptamers, while RNA anticancer aptamers and chemically modified aptamers (e.g., 2’-fluoro or locked nucleic acid modifications) were not included, which indicates that the full diversity of potential anticancer aptamer sequences may not be captured. Second, general aptamers, while providing a biologically realistic hard negative, may still contain aptamers directed against targets with indirect relevance to oncology (e.g., inflammation-or metabolism-related proteins with dual roles in cancer), which would tend to further compress the observed performance gap and should be examined through target-level curation in future work. Lastly, future experimental validation is still needed to confirm AntiCapt’s prediction ability. Furthermore, extending AntiCapt toward target-aware prediction, binding affinity estimation, and rational aptamer design could provide valuable computational support for next-generation precision oncology and accelerate the discovery of clinically relevant ACAs.

## Author Contributions

NB and GPSR collected, curated and analyzed the data. NB and NKM developed the prediction models. The results were analyzed by NB, NKM and GPSR. NB and NKM developed the front end and back end of the web server. The manuscript was prepared and finalized by NB, NKM and GPSR. The idea for this study was designed and coordinated by GPSR. All authors have examined and approved the final version of the manuscript.

## Acknowledgements

The authors would like to thank the Council of Scientific and Industrial Research (CSIR) and the Department of Biotechnology (DBT) for fellowships and financial support, and the Department of Computational Biology at IIITD, New Delhi, for the necessary infrastructure and facilities. We acknowledge <u>BioRender.com</u> for creating the Figures. The current study was supported under the grant no. BT/PR40158/BTIS/137/24/2021 provided by the Department of Biotechnology (DBT), Government of India.

## Data availability

The web server “AntiCapt” is freely available at https://webs.iiitd.edu.in/raghava/anticapt, and the dataset sequences utilized to develop the models are provided in FASTA format at https://webs.iiitd.edu.in/raghava/anticapt/dataset.php.

